# Machine learning and dengue forecasting: Comparing random forests and artificial neural networks for predicting dengue burdens at the national sub-national scale in Colombia

**DOI:** 10.1101/2020.01.14.906297

**Authors:** Naizhuo Zhao, Katia Charland, Mabel Carabali, Elaine Nsoesie, Mathieu Maher-Giroux, Erin Rees, Mengru Yuan, Cesar Garcia Balaguera, Gloria Jaramillo Ramirez, Kate Zinszer

**Affiliations:** Institute of Land Resource Management, School of Humanities and Law, Northeastern University, Shenyang, Liaoning, China; Division of Clinical Epidemiology, McGill University Health Centre, Montréal, QC, Canada; Centre de recherche en santé publique, Université de Montréal et CIUSSS du Centre-Sud-de-l’Île-de-Montréal, Montréal, Quebéc, Canada; Department of Epidemiology, Biostatistics, and Occupational Health, McGill University, Montréal, Quebec, Canada; Department of Global Health, Boston University, Boston, Massachusetts, USA; Public Health Risk Sciences Division, National Microbiology Laboratory, Public Health Agency of Canada, Saint-Hyacinthe, Québec, Canada; Faculty of Medicine, Cooperative University of Colombia, Villavicencio, Meta, Colombia; Department of Preventive and Social Medicine, University of Montreal, Montréal, Québec, Canada

**Author notes:** Corresponding author: Kate Zinszer.

## Abstract

The robust estimate and forecast capability of random forests (RF) has been widely recognized, however this ensemble machine learning method has not been widely used in mosquito-borne disease forecasting. In this study, two sets of RF models were developed for the national and departmental levels in Colombia to predict weekly dengue cases at 12-weeks ahead. A national model based on artificial neural networks (ANN) was also developed and used as a comparator to the RF models. The various predictors included historic dengue cases, satellite-derived estimates for vegetation, precipitation, and air temperature, population counts, income inequality, and education. Our RF model trained on the national data was more accurate for department-specific weekly dengue cases estimation compared to a local model trained only on the department’s data. Additionally, the forecast errors of the national RF model were smaller to those of the national ANN model and were increased with the forecast horizon increasing from one-week ahead (mean absolute error, MAE: 5.80; root mean squared error, RMSE: 11.10) to 12-weeks ahead (MAE: 13.38; RMSE: 26.82). There was considerable variation in the relative importance of predictors dependent on forecast horizon. The environmental and meteorological predictors were relatively important for short-term dengue forecast horizons while socio-demographic predictors were relevant for longer-term forecast horizons. This study showed the potential of RF in dengue forecasting with also demonstrating the feasibility of using a national model to forecast at finer spatial scales. Furthermore, sociodemographic predictors are important to include to capture longer-term trends in dengue.

**Author summary:** Dengue virus has the highest disease burden of all mosquito-borne viral diseases, infecting 390 million people annually in 128 countries. Forecasting is an important warning mechanism that can help with proactive planning and response for clinical and public health services. In this study, we compare two different machine learning approaches to dengue forecasting: random forest (RF) and neural networks (NN). National and local (departmental-level) models were compared and used to predict dengue cases in the future. The results showed that the counts of future dengue cases were more accurately estimated by RF than by NN. It was also shown that environmental and meteorological predictors were more important for forecast accuracy for shorter-term forecasts while socio-demographic predictors were more important for longer-term forecasts. Finally, the national model applied to local data was more accurate in dengue forecasting compared to the local model. This research contributes to the field of disease forecasting and highlights different considerations for future forecasting studies.

## Introduction

Dengue virus is most prevalent of the mosquito-borne viral diseases, infecting 390 million people annually in 128 countries with four different virus serotypes [1]. Rising incidence and large-scale outbreaks are largely due to inadequate living conditions, naïve populations, global trade and population mobility, climate change, and the adaptive nature of the principal mosquito vectors *Aedes aegypti* and *Aedes albopictus* [2, 3]. The direct and indirect costs of dengue are substantial and impose enormous burdens on low-and middle-income tropical countries, with a global estimate of US$8.9 billion in costs per year [4].

Human and financial costs of dengue can be alleviated when response systems, such as intervention strategies, health care services, supply chain management, receive timely warnings of future cases through forecasting models [5]. A number of dengue forecasting models have been developed and these models can be generally classified into two methodological categories: time-series and machine learning [6, 7]. The majority of existing dengue forecasting models used time-series methods and typically Autoregressive Integrated Moving Average (ARIMA), in which lagged meteorological factors (e.g. temperature and precipitation) act as covariates in conjunction with historical dengue data for one- to 12-week ahead forecasting [8–13]. Many studies reported that conventional time-series models such as ARIMA are insufficient to meet complex forecasting requirements [14–16], as multiple trends and outliers present in the time-series reduce the forecasting accuracy [17].

In the last two decades, machine learning (ML) methods have been used in many disciplines, such as geography, environment, and epidemiology, to yield meaningful findings from highly heterogeneous data. Machine learning statistical regression methods are promising approaches for disease forecasting as they facilitate the inclusion of a large number of correlated variables, enable the modeling of complex interactions between variables, and can fit complex models without strong parametric assumptions that are often untestable in traditional statistical approaches [18, 19]. Decision trees, support vector machine, artificial neural network, K-nearest neighbor, gradient boosting, and naive Bayes are frequently used ML approaches in dengue-forecasting studies [7, 20–23]. Compared to the above ML methods, random forests (RF) have shown to be more accurate in forecasting given its ability to overcome the common problem of over-fitting through the use of bootstrap aggregation [24–28].

Random forests have been used to forecast dengue risk in several countries including Costa Rica [29], Philippines [30, 31] Pakistan [32], Peru and Puerto Rico [33]. However, time or seasonal variables were not always included in the models nor were sociodemographic predictors, which have been found to improve forecast accuracy in HIV [34] and Ebola [35] epidemic models. Furthermore, dengue models, regardless of the use of the time series or ML approaches, have been developed for predicting dengue cases in individual administrative areas such in a city or a province [9–12, 20–23]. Universal dengue prediction models that are effective across different administrative regions remain absent.

Historically, Colombia is one of the countries most affected by dengue, with the *Aedes* mosquito being widely distributed throughout all departments at elevations below 2,000 meters [36, 37]. The objective of this study was to evaluate the potential of RF forecasting models at the department and national level in Colombia. We compared the accuracy of the department and national RF models to understand the feasibility of using a national model to predict dengue cases for individual departments. We also compared errors of the national RF models with those of Artificial Neural Network (ANN), another classic and widely used ML approach. Finally, we estimated the change in importance of different predictors according to forecast horizon.

## Data and methods

### Data

Various data were used to develop the forecasting models, which included: dengue cases from surveillance data, environmental indicators from remoting sensing data, and sociodemographic indicators such as population, income inequity, and education coverage (Table 1). The dengue case surveillance data were extracted from an electronic platform, SIVIGILA, created by the Colombia national surveillance program and was available at the department level. The national surveillance program receives weekly reports from all public health facilities that provide services to cases of dengue. The dengue cases reported by SIVIGILA were a mixture of probable and laboratory confirmation. Laboratory confirmation for dengue is based on a positive result from antigen, antibody, or virus detection and/or isolation [38]. Confirmation of probable cases is largely based on clinical diagnosis plus at least one serological positive immunoglobulin M test or an epidemiological link to a confirmed case 14 days prior to symptom onset.

**Table 1.**
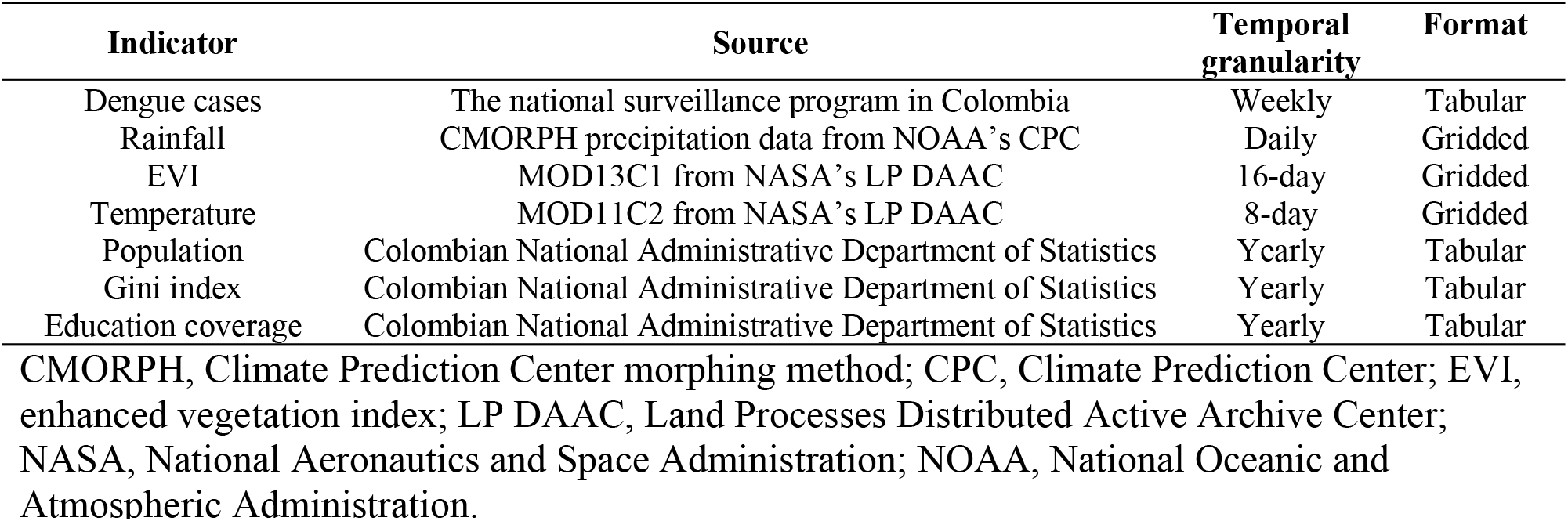
Summary of study indicators and data sources.

| Indicator | Source | Temporal granularity | Format |
| --- | --- | --- | --- |
| Dengue cases | The national surveillance program in Colombia | Weekly | Tabular |
| Rainfall | CMORPH precipitation data from NOAA's CPC | Daily | Gridded |
| EVI | MOD13C1 from NASA's LP DAAC | 16-day | Gridded |
| Temperature | MOD11C2 from NASA's LP DAAC | 8-day | Gridded |
| Population | Colombian National Administrative Department of Statistics | Yearly | Tabular |
| Gini index | Colombian National Administrative Department of Statistics | Yearly | Tabular |
| Education coverage | Colombian National Administrative Department of Statistics | Yearly | Tabular |

CMORPH, Climate Prediction Center morphing method; CPC, Climate Prediction Center; EVI, enhanced vegetation index; LP DAAC, Land Processes Distributed Active Archive Center; NASA, National Aeronautics and Space Administration; NOAA, National Oceanic and Atmospheric Administration.

Precipitation, air temperature, and land cover type have been shown to be three important determinants of *Aedes* mosquito abundance and are often used as predictors in dengue forecasting [9, 11, 21, 39]. In this study, precipitation data was obtained from the CMORPH (Climate Prediction Center morphing method) daily estimated precipitation dataset [40]. The land surface temperatures were extracted from the MODIS Terra Land Surface Temperature 8-day image products. Enhanced vegetation index (EVI) estimates were obtained from the MODIS Terra Vegetation Indices 16-Day image products. Several studies have shown that socio-demographic factors may influence dengue transmission and incidence as significantly as environmental factors [41–43]. Given this, we included population, Gini index (a measure of income inequity), and education coverage as potential predictors, which were retrieved from the Colombian National Administrative Department of Statistics. The study was approved by the Sciences and Health Ethical Committee of the University of Montreal (CERSES-19-018D), and all data were provided at the aggregate level and are publicly available.

### Random forests

Random forests (RF) is an ensemble decision tree approach [44]. A decision tree is a simple representation for classification in which each internal node corresponds to a test on an attribute, each branch represents an outcome of a test, and each leaf (i.e. terminal node) holds a class label. Decision trees can also be used for regression when the target or outcome variable is continuous. Bootstrap aggregation, commonly known as bagging, is the most distinctive technique used in RF and bagging requires training each decision tree with a randomly selected subsample of the entire training datasets.

### Data preprocessing

To ensure a consistent temporal granularity with the outcome variable, the daily precipitation data were aggregated to a weekly frequency. The 8-day land surface temperature and the 16-day EVI data were resampled to a weekly frequency using a spline interpolation [45]. We assigned a given department the same population, Gini index, and education coverage values for all weeks within the same calendar year.

The archipelago of San Andrés, Providencia, and Santa Catalina (commonly known as *San Andrés y Providencia*) is a department consisting of two island groups and 775 km away from mainland Colombia. Due to the frequent cloud contamination over the small island areas, it was not possible to have high-quality MODIS images products for weekly temperature or EVI value estimation. Vaupés department had only 30 confirmed dengue cases scattered in 24 weeks during 2014 to 2018. Thus, the departments of San Andrés y Providencia and Vaupés were excluded from this study, and data from the other 30 departments were used to train our models.

Weekly dengue data from 2014-2017 was used to train the RF models and the data from 2018 was used to evaluate the models. To simulate ‘real life’ forecasting, we did not include the 2018 data for the socio-demographic variables given that they are only produced annually whereas the remote sensing data are more readily available. Exponential smoothing approach based on historical (2010-2017) time-series data to estimate the values for 2018.

### Development of RF models

We first developed RF models for each department (referred to as local level). Let the “current” week be *k* and the number of confirmed dengue cases be *y*. Referring to the RF streamflow forecasting model developed by Papacharalampous and Tyralis [46], we used the numbers of current and previous 11 weeks dengue cases (i.e. *y_k_*, *y_k-1_*, …, *y_k-10,_ y_k-11_*) of a department to predict one-week ahead dengue cases (i.e. *y_k+1_*) for each department. The current and previous 11 weeks of rainfall, land surface temperature, EVI, population, Gini index, and education coverage were also included as predictors. These values were selected as previous studies demonstrated that the optimal lags of meteorological variables used for dengue forecasting are usually not larger than 12 weeks [47–52]. In addition, the ordinal number of the forecast week (1–52 for the year of 2015, 2016, 2017, and 2018 and 1–53 for 2014) as well as year (2014–2018) were treated as two predictor variables to account for seasonality and long-term changing trend of dengue occurrence [53,54].

We then developed RF models at the national scale. To train a national-scale RF model for forecasting *n*-week ahead dengue cases (where *n* ≤12), we used the same predictor and target variables as those used in the local *n*-week ahead forecasting models. The difference between the local and the national models was that the local *n*-week ahead models were trained using 209-*n* (209 =53+52+52+52) samples while the national model was trained using 6270-30*n* [i.e. (209-n) ×30] samples.

### Model evaluation

Model accuracy was evaluated and compared by two metrics: mean absolute error (MAE) and root mean squared error (RMSE). The MAEs and RMSEs reported in this study were calculated by the actual and the predicted numbers of dengue cases for the 52 weeks in 2018. The accuracy comparison was performed at the local (department) and national scales. When the comparison for an *n*-week ahead prediction was conducted at the national scale, the predicted numbers of dengue cases by the 30 local RF models were additively combined and compared with the actual national values to calculate one MAE and one RMSE. When the comparison was implemented at the local scale, the national RF model was applied to each one of the 30 departments and then the predicted values were compared with the actual numbers of dengue cases to compute 30 individual MAEs and 30 individual RMSEs.

Artificial Neural Network (ANN) is an early ML approach and has been previously used to predict dengue cases [7, 20, 21, 23]. We developed ANN models at the national scale and compared their prediction accuracy with that of the RF models. The ANN was composed of one input layer, one hidden layer, and one output layer. The ANN models had the same 53 predictor variables as the RF models, resulting in 53 neurons in the input layer and one neuron in the output layer. The number of neurons in the hidden layer was determined by iterative attempts until the prediction accuracy cannot be further improved [55]. In this study, the optimal number of neurons in hidden layer varied by forecasting horizon and ranged between 38 to 50.

Percentage of increased mean squared error (%IncMSE) is a robust and informative indicator to quantitatively evaluate the importance of predictor variables in a random forests model [56]. Percentage of increased mean squared error indicates the increase in the mean squared error (MSE) of prediction as a result of an independent variable being randomly shuffled while maintaining the other independent variables as unchanged [44]. A larger %IncMSE of a predictor variable suggests greater importance of the variable on the model’s overall forecast accuracy and the %IncMSE was calculated for each predictor in each RF model.

## Results

An exceptionally large dengue outbreak occurred in Colombia during the study period. The counts of confirmed dengue cases reached more than 2,500 per week by the end of 2015 and the outbreak ended mid-year in 2016. Following this outbreak, the yearly dengue case peaks were drastically reduced in 2016 and 2017 but began increasing again in 2018 (Fig1).

**Fig 1.**
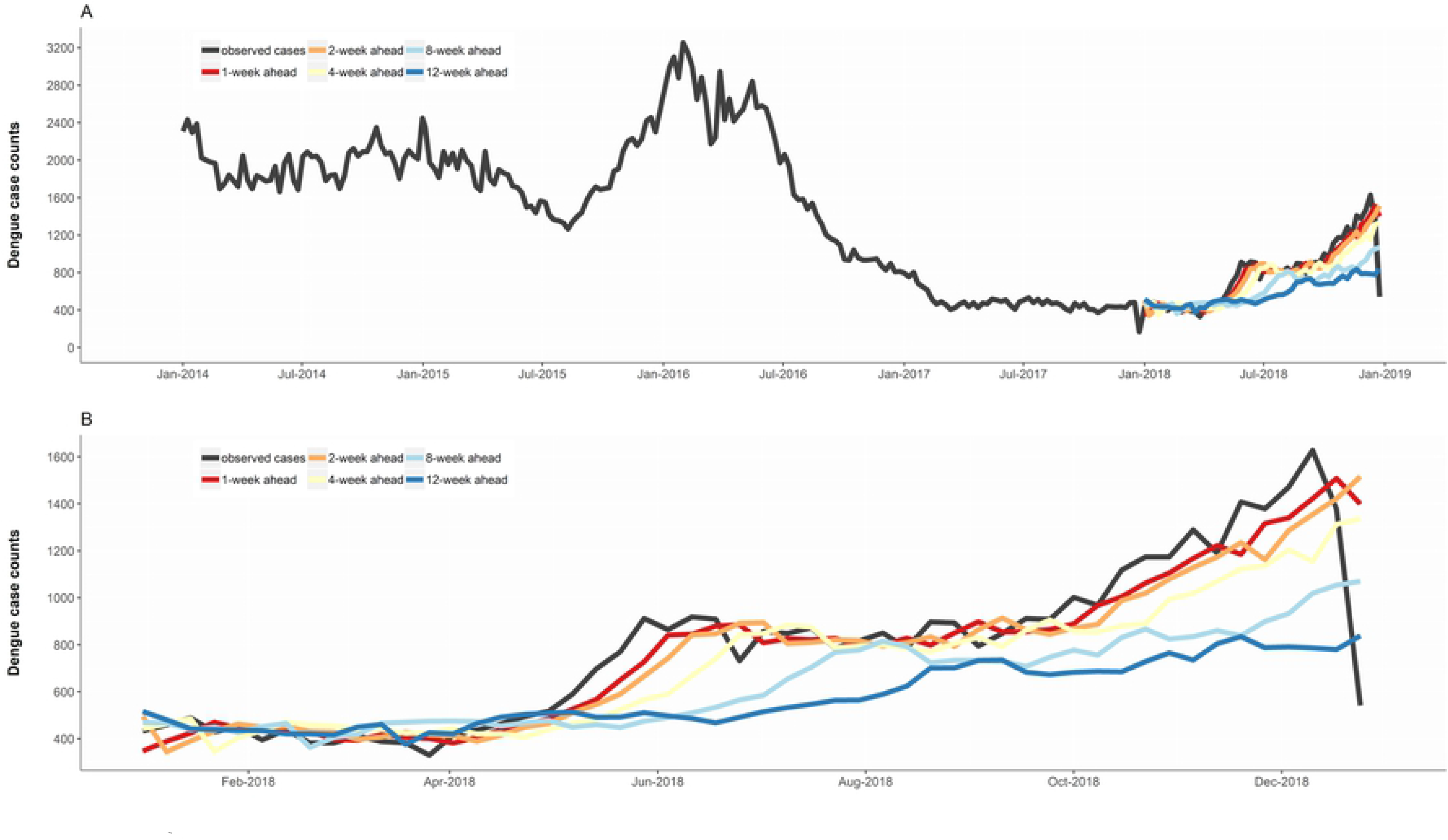
The weekly total counts of confirmed dengue cases over Colombia for 2014-2018 (A) and the predicted counts of dengue cases by the national one-, two-, four-, eight-, and twelve-week ahead models for 2018 (B). See Fig S1 for the predicted counts of dengue cases for all week ahead models.

For any of the n-week ahead (n≤12) forecasts, the performance of the national model was better than that of the local model, demonstrated by the smaller overall MAE and RMSE (Table 2). Moreover, in most cases, a department’s dengue cases were more accurately predicted by the national model than the local model (Fig 2). The errors of the national random forests model were mainly derived from under-estimation of cases which coincided with dramatic increases in cases towards the end of 2018. As expected, the under-estimation was more pronounced when predictions were made over a longer time period.

**Table 2.**
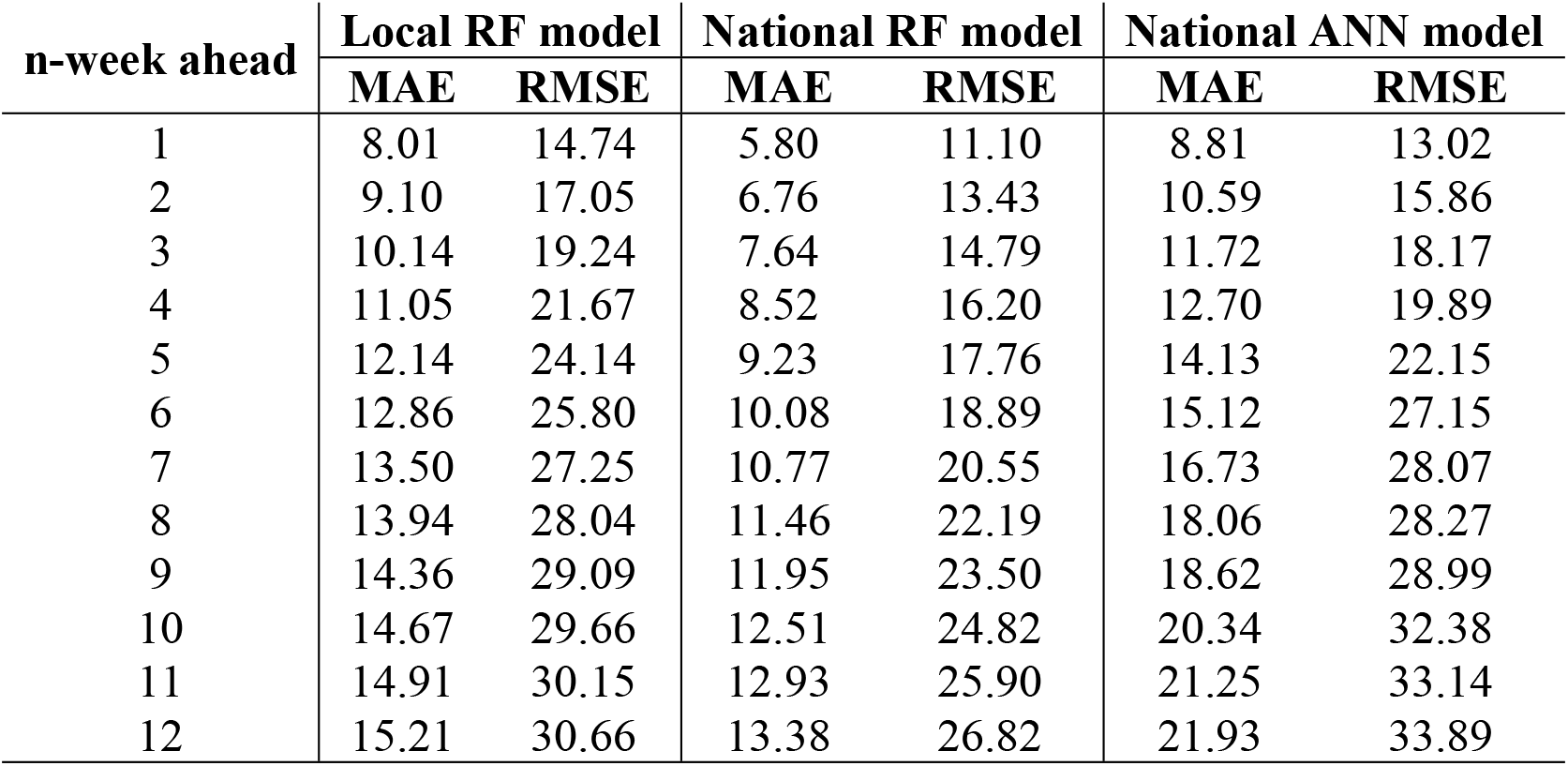
Comparison of accuracy between the local and the national models. MAE, mean absolute error; RMSE, root mean squared error; RF, random forests; ANN, artificial neural network.

**Fig 2.**
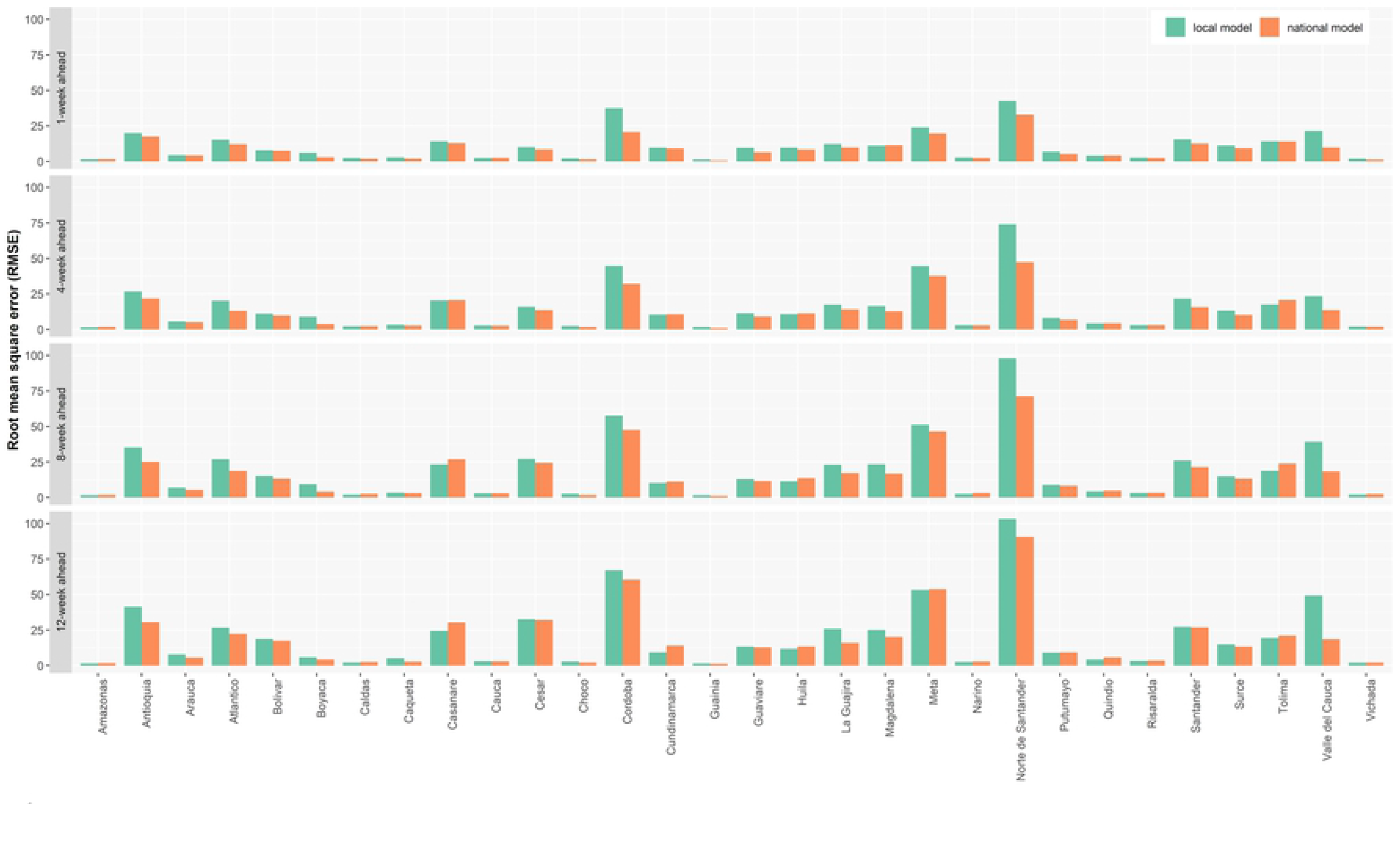
Accuracy comparison between the local and the national random forests models at the department scale for the one-week ahead, four-week ahead, eight-week ahead, and twelve-week ahead predictions with RMSE for 2018. See Fig S2-S4 in the supporting information on MAE and for all week ahead models.

The overall RMSE of the ANN model developed at the national scale was smaller than that of the local RF model at forecasting horizons of 5 weeks or less (Table 2). The RMSE grew for the ANN model with longer forecasting horizons compared to the local RF model. The MAE of the ANN model was consistently larger than that of the local RF model for each forecasting horizon. The RMSE and MAE of the national RF model were smaller than those of the national ANN model at any forecasting horizon.

The relative importance of different predictor variables in the national RF model was varied (Table 3). Firstly, “current” and “near current” past dengue data were extremely important in predicting occurrence of dengue in the near future (e.g. one- to three-weeks ahead). However, with the predicted week increasingly further away from the “current” week, the importance of historical dengue data decreased while the “current” week of dengue cases remained one of the top three most important predictors in predicting the future dengue cases. Secondly, the environmental (EVI) and the meteorological predictors (rainfall and temperature) were more important than the socio-demographic predictors when dengue cases were predicted in the near future (one- to three-weeks ahead). Yet, with the predicted week increasingly far away from the “current” week, the three socio-demographic covariates (education, population, and Gini index) became increasingly important. Finally, the week predictor, which accounted for the seasonal pattern of dengue, was important across all forecasting horizons but relatively smaller in importance with smaller forecasting horizons (i.e. *n* ≤4)

**Table 3.** The top ten most important predictor variables for predicting dengue cases in the national models, ordered from the largest to the smallest %IncMSEs.

| Rank | 1 | 2 | 3 | 4 | 5 | 6 | 7 | 8 | 9 | 10 |
| --- | --- | --- | --- | --- | --- | --- | --- | --- | --- | --- |
| 1-week ahead | Dengue<sub>k</sub> | Dengue<sub>k-1</sub> | Dengue<sub>k-2</sub> | Dengue<sub>k-3</sub> | Week | Dengue<sub>k-4</sub> | EVI<sub>k-11</sub> | Temperature<sub>k-11</sub> | EVI<sub>k-10</sub> | EVI<sub>k-8</sub> |
| 2-week ahead | Dengue<sub>k</sub> | Dengue<sub>k-1</sub> | Week | Dengue<sub>k-2</sub> | Dengue<sub>k-3</sub> | Temperature<sub>k-11</sub> | Dengue<sub>k-4</sub> | EVI<sub>k-7</sub> | EVI<sub>k-5</sub> | EVI<sub>k-8</sub> |
| 3-week ahead | Dengue<sub>k</sub> | Dengue<sub>k-1</sub> | Week | Dengue<sub>k-2</sub> | EVI<sub>k-8</sub> | EVI<sub>k-10</sub> | Temperature<sub>k-10</sub> | Education | Dengue<sub>k-3</sub> | Dengue<sub>k-4</sub> |
| 4-week ahead | Dengue<sub>k</sub> | Week | Dengue<sub>k-1</sub> | Education | Dengue<sub>k-2</sub> | Temperature<sub>k-9</sub> | EVI<sub>k-8</sub> | Temperature<sub>k-11</sub> | EVI<sub>k-7</sub> | Dengue<sub>k-3</sub> |
| 5-week ahead | Dengue<sub>k</sub> | Week | Dengue<sub>k-1</sub> | Education | Dengue<sub>k-2</sub> | EVI<sub>k-10</sub> | Temperature<sub>k-8</sub> | Temperature<sub>k</sub> | Gini | EVI<sub>k-9</sub> |
| 6-week ahead | Dengue<sub>k</sub> | Week | Dengue<sub>k-1</sub> | Education | Population | Year | Dengue<sub>k-2</sub> | EVI<sub>k-8</sub> | EVI<sub>k-9</sub> | EVI<sub>k-10</sub> |
| 7-week ahead | Dengue<sub>k</sub> | Week | Education | Dengue<sub>k-1</sub> | Year | Dengue<sub>k-2</sub> | Population | Gini | EVI<sub>k-10</sub> | EVI<sub>k-9</sub> |
| 8-week ahead | Dengue<sub>k</sub> | Week | Population | Education | Dengue<sub>k-1</sub> | Year | Temperature<sub>k-11</sub> | Temperature<sub>k-5</sub> | Dengue<sub>k-2</sub> | Gini |
| 9-week ahead | Dengue<sub>k</sub> | Week | Population | Education | Year | Dengue<sub>k-1</sub> | Temperature<sub>k-11</sub> | Dengue<sub>k-11</sub> | Gini | Temperature<sub>k-3</sub> |
| 10-week ahead | Dengue<sub>k</sub> | Week | Year | Education | Population | Dengue<sub>k-1</sub> | Gini | Dengue<sub>k-11</sub> | Temperature<sub>k-4</sub> | Dengue<sub>k-2</sub> |
| 11-week ahead | Year | Week | Dengue<sub>k</sub> | Population | Education | Gini | Dengue<sub>k-1</sub> | Temperature<sub>k-11</sub> | Dengue<sub>k-10</sub> | Temperature<sub>k-4</sub> |
| 12-week ahead | Population | Year | Dengue<sub>k</sub> | Week | Education | Gini | Dengue<sub>k-11</sub> | Dengue<sub>k-1</sub> | Dengue<sub>k-10</sub> | Temperature<sub>k-10</sub> |

Dengue indicates historical dengue cases and EVI denotes enhanced vegetation index. %IncMSE, percentage of increased mean squared error.

## Discussion

In the current study, we developed a national model to predict counts of dengue cases across different departments of Colombia and found that for the majority of departments, the national model more accurately forecasted future dengue cases at the department level compared to the local model. This result indicates the similarity in importance of dengue drivers across different administrative regions of Colombia. Random forests is an unsupervised tree-based regression approach requiring a relatively large training sample for the repeated splitting of the dataset into separate branches, and thus the national model trained by a larger dataset had higher prediction accuracy compared to the local models. The national and the local models performed poorly in departments of Guainía and Vichada. The small population and consequently, the low counts of dengue cases resulted in the relatively large errors in the two departments.

We found that the meteorological and environmental variables were more important for prediction accuracy at smaller forecasting horizons compared to the socio-demographic variables, with socio-demographics being more important at larger forecasting horizons. This is likely due to the influence of meteorological and environmental conditions on *Aedes* mosquitoes and the lag effects are usually between 1 to 4 weeks for temperature and precipitation [57–59]. Poor quality housing and sanitation management with high population density are key risk factors for dengue transmission [60, 61], and are closely related to education and poverty [62, 63]. These results demonstrate the complimentary nature of these different groups of predictor variables and the importance of their inclusion in dengue forecasting models.

We used ANN models as comparators to our RF models. Artificial Neural Networks are brain-inspired systems that are intended to imitate the way that human learn. Theoretically, more complex correlations between predictor and target variables can be discerned by deeper (i.e. more hidden layers) networks [64]. However, ANN cannot handle the problem of vanishing gradient which results in the failure of improving accuracy of ANN models by adding more hidden layers. Additionally, it is easy for ANN to suffer from over-fitting which leads to a network developed by a training dataset failing to predict the other observations accurately. In this study, the number of neurons in the hidden layer was required to be changed with each forecast horizon, demonstrating the poor universality of the ANN models. By contrast, RF solves the problem of over-fitting with the use of bootstrap aggregation. Hyperparameters (e.g. the number of decision trees) in RF are easy to be set and the RF models showed better universality for different forecast horizons.

Despite the strengths of our study, an important limitation with our RF approach is that the considerable dependence on the current week of dengue leads the model to generate lags for forecasting rapid changes in dengue. Including a predictor of mosquito abundance from an entomological surveillance program may reduce such time lag errors [65]. However, this type of data is often difficult to obtain at the national level with sufficient temporal and spatial granularity. Additionally, RF, as a non-parametric black-box approach, cannot intuitively display quantitative relationships between the count of dengue cases and the heterogeneous predictor variables, although it is able to more flexibly and accurately model the possibly complex non-linear and non-additive relationships among the variables. A more severe limitation of the RF model is the fact that RF cannot obtain values beyond the range of the variable in the training dataset. If an unprecedented dengue outbreak occurred in future, under-estimations will occur inevitably using the RF approach.

## Conclusions

This study highlights the potential of RF for dengue forecasting and also demonstrates the benefits of including socio-demographic predictors. Our findings also found that a national model, on average, performed better compared to the local models. Future studies should consider the inclusion of other arboviruses as predictors, such as chikungunya and Zika as well as examine the importance of other socio-economic factors. In addition, other promising ML methods should be tested including recurrent neural networks, which inherently account for time, have the ability to deal with a vanishing gradient, and are able to capture complicated non-linear and non-additive relationships between predictor and target variables [66].

## Support Information Legends

Fig S1. The weekly total counts of confirmed dengue cases over Colombia for 2014-2018 (A) and the predicted counts of dengue cases by the national model for one to twelve-week ahead for 2018 (B).

Fig S2. Accuracy comparison between the local and the national random forests models at the department scale for the one-week ahead, four-week ahead, eight-week ahead, and twelve-week ahead predictions with MAE for 2018.

Fig S3. Accuracy comparison between the local and the national random forests models at the department scale for one to twelve-week ahead predictions with RSME for 2018.

Fig S4. Accuracy comparison between the local and the national random forests models at the department scale for one to twelve-week ahead predictions with MAE for 2018.

